# High-accuracy protein complex structure modeling based on sequence-derived structure complementarity

**DOI:** 10.1101/2025.03.26.645390

**Authors:** Minghua Hou, Yuhao Xia, Pengcheng Wang, Zexin lv, Dongliang Hou, Xiaogen Zhou, Jianyang Zeng, Guijun Zhang

## Abstract

In living organisms, proteins perform key functions required for life activities by interacting to form complexes. Determining the protein complex structure is crucial for understanding and mastering biological functions. Although AlphaFold2 has made a revolutionary breakthrough in predicting protein monomeric structures, accurately capturing inter-chain interaction signals and modeling the structures of protein complexes remain a formidable challenge. In this work, we report DeepSCFold, a pipeline for improving protein complex structure modeling. DeepSCFold uses sequence-based deep learning models to predict protein-protein structural similarity and interaction probability, providing a foundation for identifying interaction partners and constructing deep paired multiple-sequence alignments (MSAs) for protein complex structure prediction. Benchmark results show that DeepSCFold significantly increases the accuracy of protein complex structure prediction compared with state-of-the-art methods. For multimer targets from CASP15, DeepSCFold achieves an improvement of 11.6% and 10.3% in TM-score compared to AlphaFold-Multimer and AlphaFold3, respectively. Furthermore, when applied to antibody-antigen complexes from the SAbDab database, DeepSCFold enhances the prediction success rate for antibody-antigen binding interfaces by 24.7% and 12.4% over AlphaFold-Multimer and AlphaFold3, respectively. These results demonstrate that DeepSCFold effectively captures intrinsic and conserved protein-protein interaction patterns through sequence-derived structure-aware information, rather than relying solely on sequence-level co-evolutionary signals.

## Introduction

Interacting proteins play pivotal roles in cellular processes by forming functional multi-protein complexes (i.e., multimers or assemblies) that are essential for executing biological processes such as signal transduction, transport, and metabolism^1–4^. To fully understand these functions, determining the structures of these complexes is crucial. However, this often remains a challenge for existing experimental methods, such as X-ray crystallography, nuclear magnetic resonance (NMR), and cryo-electron microscopy (cryo-EM)^5^. Consequently, computational methods for obtaining complex structures have become an indispensable and important complement to experimental techniques in structural biology. Nevertheless, predicting the quaternary structure of a protein complex is significantly more challenging than predicting the tertiary structure of a single protein monomer, as it necessitates the accurate modeling of both intra-chain and inter-chain residue-residue interactions among multiple protein chains^6^.

Traditionally, the prediction of protein complex structures relies on template-based homology modeling or docking-based prediction methods^6,7^. While template-based homology modeling is effective when high-quality templates are available, its applicability is often limited by the difficulty in obtaining suitable templates for most target complexes^8^. In protein-protein docking methods, monomer structures are assembled into a complex based on the "lock-and-key"^9^ and "induced fit"^10^ principles, as exemplified by tools such as ZDOCK^11–13^, HADDOCK^14–16^, and HDOCK^17–19^. The docking process aims to identify the optimal binding mode that satisfies the spatial shape complementarity through energy minimization. However, docking methods still face challenges in predicting complex structures due to the complexity of conformational sampling, the inaccuracy of energy functions, and the inherent flexibility of proteins in the interface regions^20,21^.

With the remarkable advancements achieved by AlphaFold2^22^ in protein monomer structure prediction, deep learning methods have increasingly been applied to predict protein multimer structures, including AlphaFold-Multimer^23^, DMFold-Multimer^24,25^, MULTICOM3^6^, and the recently released AlphaFold3^26^. Among these, AlphaFold-Multimer, an extension of AlphaFold2 specifically tailored for protein multimer structure prediction, has significantly improved the accuracy of complex predictions. Nevertheless, the accuracy of multimer structure predictions using AlphaFold-Multimer remains considerably lower than that of AlphaFold2 for monomer structures^6^. To address this limitation, the research community has actively developed strategies based on AlphaFold-Multimer to further enhance performance. These strategies include extensive sampling through variations in multiple sequence alignments (MSAs) construction, the use of multiple seeds, increased recycling, and extensive network dropout^27^. Results from CASP15 highlight the effectiveness of these methods, demonstrating superior performance in predicting protein multimers compared to the baseline AlphaFold-Multimer (i.e., the NBIS-AF2-standard group)^28^. Notable examples include the Zheng group^25^, Venclovas group^29^, Wallner group^30^, Yang-Multimer group^31^ and MULTICOM_human group^6^ (i.e., MULTICOM3), etc. Among these advancements, the quality of MSAs plays a critical role in structure prediction, as their implicit co-evolutionary information is often essential for locating an approximate global minimum in the protein conformation space^28,32^. This importance is further amplified in the context of protein complexes, where accurately capturing the binding modes of protein chains may significantly benefit from the paired MSAs of the complex.

In protein complex structure prediction, monomeric MSAs derived from individual chains are systematically paired across different chains to generate comprehensive paired MSAs. This pairing strategy enables the identification of inter-chain co-evolutionary signals between interacting partners, providing valuable insights into the dynamic behavior and stability of molecular interactions within the protein complex. However, popular sequence search tools such as HHblits^33,34^, Jackhammer^35^, and MMseqs^36,37^ are primarily designed for constructing monomeric MSAs and cannot be directly applied to paired MSAs construction. This limitation may compromise the accuracy and generality of protein complex structure predictions, particularly for tightly intertwined complexes or highly flexible interactions, such as those observed in antibody-antigen systems^38^. Recently, several methods have been proposed to construct paired MSAs, aiming to enhance the ability to capture inter-chain interactions. Examples include DeepMSA2^24^, MULTICOM^36^, DiffPALM^38^, ESMPair^39^. DeepMSA2 improves monomeric MSAs and constructs paired MSAs by performing iterative alignment searches across genomic and metagenomic sequence databases, followed by filtering using AlphaFold2/AlphaFold-Multimer. MULTICOM3 generates a diverse set of paired MSAs by concatenating the MSAs of subunits, leveraging potential protein-protein interactions extracted from multiple sources. ESMPair ranks monomeric MSAs using ESM-MSA-1b^40^ and integrates species information to construct paired MSAs. DiffPALM employs an MSA transformer to estimate amino acid probabilities, creating a permutation matrix to pair protein sequences. While these methods effectively capture inter-chain co-evolutionary information through paired MSAs construction based on sequence similarity or species-related information, they may face limitations when applied to complexes lacking clear co-evolutionary signals at the sequence level. For instance, virus-host and antibody-antigen systems often do not exhibit inter-chain co-evolution, as identifying orthologs between host and pathogenic proteins is challenging due to the absence of species overlap. In such a case, leveraging deep learning architectures to directly extract spatial conformational complementarity features from each individual monomeric protein sequences represents a transformative approach, offering both critical structural insights and significant potential for enhancing the precision of protein complex structure prediction^41^.

Protein structures are generally more functionally conserved than their corresponding sequences due to their direct involvement in mediating biological processes. This evolutionary conservation is particularly evident at the structural level of protein-protein interactions (PPIs), where interaction interfaces tend to be more conserved than sequence motifs. Extensive experimental evidence suggests that the repertoire of protein interaction modes in nature is remarkably limited^42,43^, with similar structural binding patterns observed across diverse PPIs^44,45^. Motivated by these fundamental observations, we present DeepSCFold, a novel deep learning framework that systematically captures structural complementarity^46,47^ between protein chains through the integration of protein sequence semantic embeddings as well as physicochemical and statistical features. Our method enables comprehensive exploration of protein binding modes and accurate prediction of complex quaternary structures. Benchmark evaluations on the CASP15 protein complex dataset demonstrate that DeepSCFold outperforms existing state-of-the-art methods in both global and local interface accuracy. To further validate the robustness of our method, we conducted specialized evaluations using antibody-antigen complexes from the SAbDab^48^ database, focusing on challenging cases that lack detectable inter-chain co-evolution signals. Our results demonstrate that structural complementarity-based paired MSAs can effectively compensate for the absence of co-evolutionary information by providing reliable inter-chain interaction signals.

## Results

### Overview of the method

DeepSCFold is a computational protocol specifically developed for high-accuracy prediction of protein complex structures. The method constructs paired multiple sequence alignments (pMSAs) by integrating two key components: (1) assessing structural similarity between monomeric query sequences and their corresponding homologs within individual MSAs, and (2) identifying potential interaction patterns among sequences across distinct monomeric MSAs. This dual-strategy approach enables the systematic generation of high-quality pMSAs, which serve as the foundation for accurate protein complex modeling. At the core of DeepSCFold’s paired-MSA construction are two advanced sequence-based deep learning models: one predicts protein-protein structural similarity (pSS-score) purely from sequence information, while the other estimates interaction probability (pIA-score) based solely on sequence-level features. These models enable the inference of structural and interaction properties without relying on prior structural knowledge, making DeepSCFold uniquely capable of modeling complex interactions from sequence data alone. **Fig. 1** shows an overview of the DeepSCFold protocol. Starting from the input protein complex sequences, DeepSCFold first generates monomeric multiple sequence alignments (MSAs) from multiple sequence databases (UniRef30^49^, UniRef90^50^, UniProt^50^, BFD^51,52^, MGnify^53–55^, and the ColabFold DB^56^). Then, the predicted pSS-score, which quantifies the structural similarity between the input sequence and its corresponding homologs in the monomeric MSAs, was employed as a complementary metric to traditional sequence similarity, thereby enhancing the ranking and selection process of monomeric MSAs. Subsequently, the developed deep learning model predicts the pIA-scores for each potential pair of sequence homologs derived from distinct subunit MSAs. These interaction probabilities are then utilized to systematically concatenate monomeric homologs and construct paired MSAs, enabling the identification of biologically relevant interaction patterns. Additionally, we integrate multi-source biological information, including species annotations, UniProt accession numbers, and experimentally determined protein complexes from the Protein Data Bank (PDB), to construct additional paired MSAs with enhanced biological relevance. Finally, DeepSCFold uses the series of paired MSAs constructed above to perform complex structure predictions through AlphaFold-Multimer. The top-1 model is selected based on our in-house complex model quality assessment method DeepUMQA-X and then used as the input template of AlphaFold-Multimer for one iteration to generate the final output structure.

**Fig. 1.**
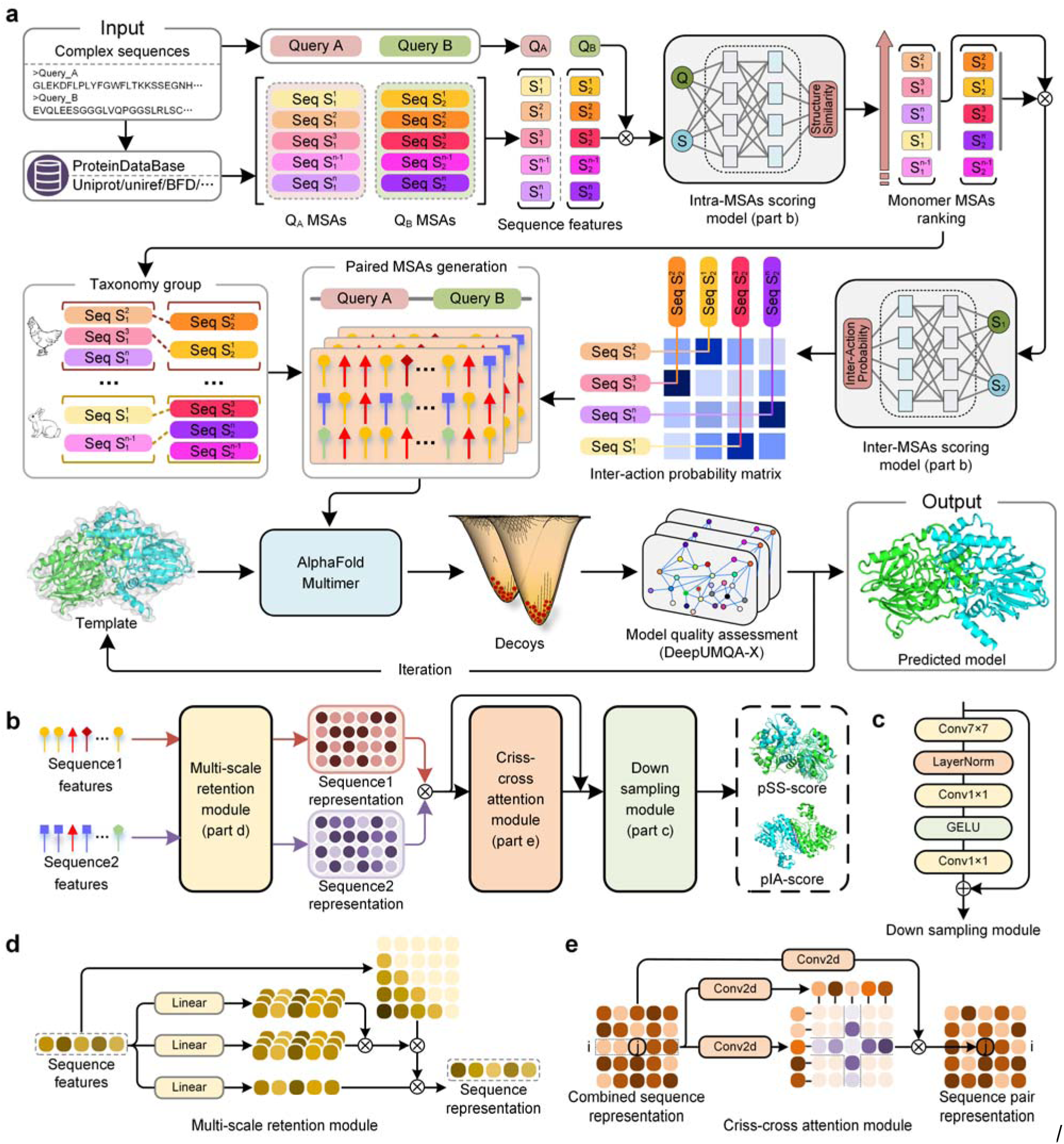
Pipeline of DeepSCFold. **a.** DeepSCFold takes multiple query sequences of a protein complex as input, with each component sequence being independently searched against protein databases to generate monomeric multiple sequence alignments (MSAs). A deep learning model is designed to assess structural similarity between each sequence homolog in the MSA and the query sequence, enabling the ranking and selection of appropriate homologs. The pairing of homologs from different monomeric MSAs is accomplished through two distinct strategies: one leveraging species information or manually curated annotations, and another utilizing a probability matrix derived from an interaction probability prediction model that scores potential homolog pairs. These paired MSAs are subsequently processed by AlphaFold-Multimer to generate potential complex structures. The resulting models undergo quality assessment, with the top-ranked model serving as a template for a single iteration of refinement, ultimately yielding the final predicted structure of the protein complex. **b.** Deep learning model architecture: The features of a pair of sequences are input into a multi-scale retention module (**d**) to generate sequence representations, which are then fed into a criss-cross attention module (**e**) to generate paired representations, followed by a down sample module (**c**) to generate the final predicted scores (pSS or pIA).

### Improvements of protein complex structure prediction by DeepSCFold

To evaluate the performance of DeepSCFold in predicting protein complex structures, we systematically tested the method on a benchmark set of multimeric targets from the CASP15 competition. For each target, complex models were generated using protein sequence databases available up to May 2022, ensuring a temporally unbiased assessment of our method’s predictive capabilities. The predictions were then compared with those from several state-of-the-art methods, including AlphaFold3, Yang-Multimer, MULTICOM, and NBIS-AF2-multimer. The prediction results of AlphaFold3 models were generated using its online server (https://golgi.sandbox.google.com/), while the last three methods were retrieved from the CASP15 official website. To assess the accuracy of the predicted models, we employed two complementary metrics: TM-score, which quantifies the global topological similarity to the experimentally determined structure, and DockQ, which specifically evaluates the quality of protein-protein docking interfaces. A detailed description of these metrics is provided in **Supplementary Note 1**.

**Fig. 2a** shows the TM-score and DockQ of the predicted multimer protein models with respect to the experimental structures. The detailed evaluation results are listed in **Supplementary Table 1**. DeepSCFold demonstrates superior performance in protein complex structure prediction, achieving an average TM-score of 0.87 and DockQ score of 0.59. This represents a significant improvement over current state-of-the-art methods, including AlphaFold3 (TM-score = 0.78, DockQ = 0.49), Yang-Multimer (TM-score = 0.82, DockQ = 0.51), MULTICOM (TM-score = 0.82, DockQ = 0.49), and NBIS-AF2-multimer (TM-score = 0.78, DockQ = 0.43). The consistent enhancement across both evaluation metrics highlights DeepSCFold’s advanced capabilities in modeling protein complexes with greater accuracy and reliability. These findings indicate that the integration of sequence-derived structural complementarity in DeepSCFold likely enhances its ability to generate protein multimer models with improved global topological accuracy and biologically meaningful docking modes, as evidenced by its higher TM-score and DockQ. Given that NBIS-AF2-multimer serves as the implementation of AlphaFold-Multimer in CASP15, we performed a direct comparative analysis between DeepSCFold and NBIS-AF2-multimer to evaluate the effectiveness of our enhanced input strategy in improving the accuracy of multimer structure predictions. **Fig. 2b** provides a detailed comparison of prediction accuracy between DeepSCFold and NBIS-AF2-multimer for each individual target. The results demonstrate that DeepSCFold outperforms NBIS-AF2-multimer, achieving superior TM-scores in 92% of the test cases and higher DockQ scores in 88% of the cases. To provide a more detailed evaluation, we categorized the results based on different performance thresholds for TM-score (0.5 and 0.9) and DockQ (0.23, 0.49, and 0.8), allowing for a nuanced comparison across various quality levels. Several cases show notable improvements in both metrics, with models improving from unreliable to good or high-quality levels, such as intertwined complexes (e.g., T1161), nanobody-antigen complexes (e.g., H1144), and antibody-antigen complexes (e.g., H1166).

**Fig. 2.**
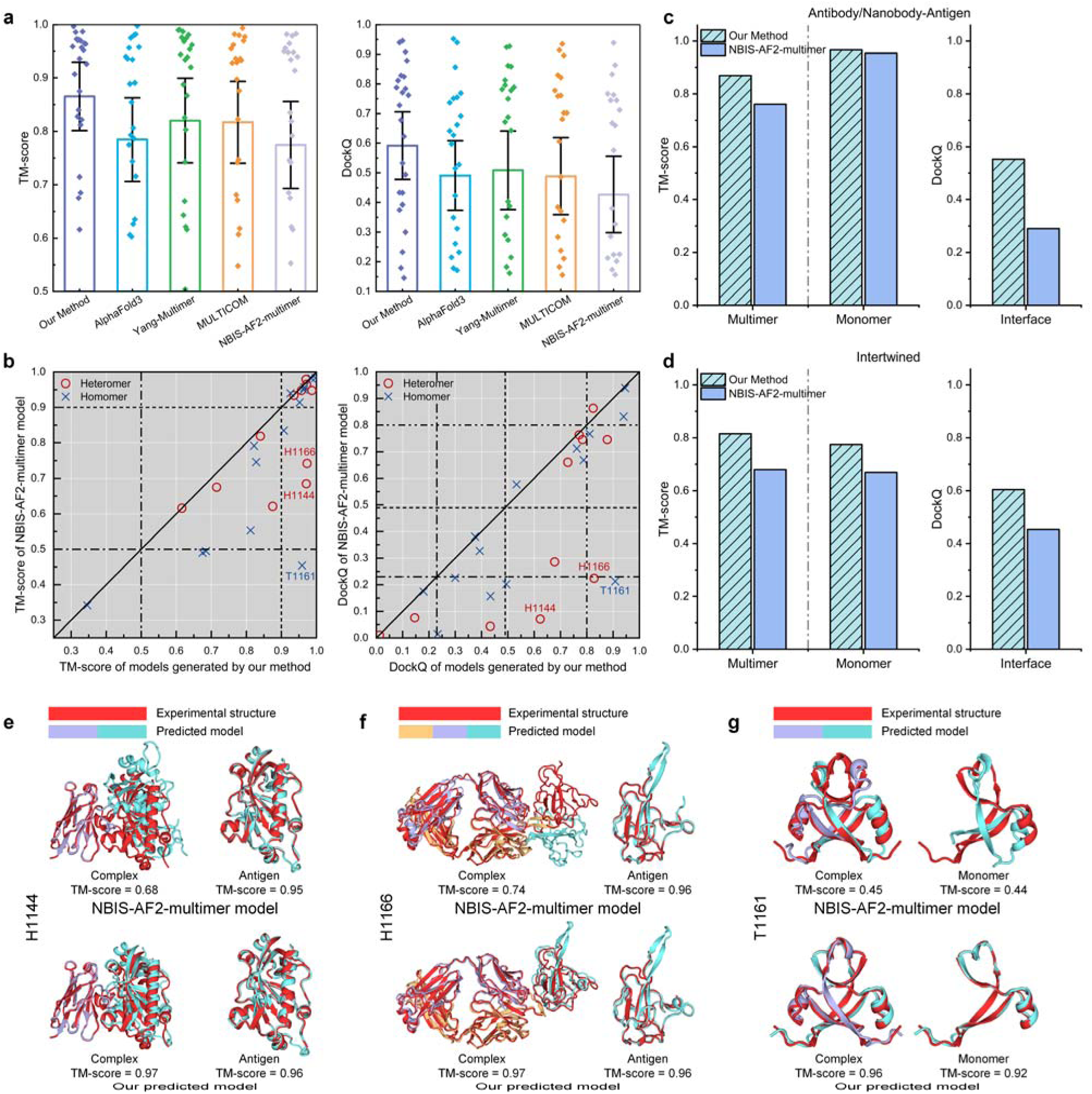
Performance of DeepSCFold in protein complex structure prediction on the CASP15 dataset. **a.** Average TM-scores and DockQ scores of the complex models by DeepSCFold, Yang-Multimer, MULTICOM, NBIS-AF2-multimer, and AlphaFold3. Error bars are 95% confidence intervals. **b.** Head-to-head comparison of the TM-scores and DockQ scores on each test case between DeepSCFold and NBIS-AF2-multimer, with three representative cases labeled in the figure and detailed analysis provided in (**e**), (**f**) and (**g**). **c-d.** Performance comparison for antibody/nanobody-antigen complexes (**c**) and intertwined complexes (**d**), showing average TM-scores of multimers and monomers as well as interface DockQ scores. **e-g.** Examples of the multimer structures built by DeepSCFold and NBIS-AF2-multimer for nanobody-antigen complex H1144 (**e**), antibody-antigen complex H1166 (**f**) and intertwined complex T1161 (**g**), with experimental structures shown in red and the monomers of the predicted models colored differently. The H1144 and H1166 complexes are aligned based on the antibody chains.

To further elucidate the underlying mechanisms of these improvements, we conducted a in-depth analysis of these complex types, as illustrated in **Fig. 2c** and **Fig. 2d**. This focused investigation provides valuable insights into the specific structural features and interactions contributing to the enhanced predictive performance. **Fig. 2c** shows a comparison of the results for the antibody/nanobody-antigen complex, showing the average multimer TM-score, monomer TM-score, and interface DockQ. Both methods demonstrate robust performance in monomer prediction, with DeepSCFold attaining a monomer TM-score of 0.97 and NBIS-AF2-multimer achieving 0.95. However, the critical distinction lies in their interface prediction capabilities. DeepSCFold achieves a substantially higher interface DockQ score of 0.55, outperforming NBIS-AF2-multimer’s score of 0.29. This marked enhancement in interface prediction accuracy directly contributes to DeepSCFold’s superior multimer TM-score of 0.87, compared to 0.76 for NBIS-AF2-multimer, highlighting the importance of precise interface modeling in multimer structure prediction. As a representative case study, in **Fig. 2e** we present an example from H1144, which is a nanobody-antigen complex containing two protein chains with 341 residues in total. Although NBIS-AF2-multimer generates a high-quality monomer model with a TM-score of 0.95, the quaternary orientation of the complex is completely wrong resulting in a poor complex TM-score of 0.68. In contrast, our method provides more accurate interchain signals, leading to a significantly improved complex model with a TM-score of 0.97. **Fig. 2f** shows another illustrative case from H1166, which is an antibody-antigen complex with 577 residues in total. NBIS-AF2-multimer achieves a high monomer TM-score of 0.96, but the complex TM-score is only 0.74, whereas our model achieves 0.97. Antibody/nanobody-antigen complexes are inherently cross-species systems, which lack intra-species binding signals in their respective subunit sequence alignments (MSAs). Consequently, the MSAs generated by the NBIS-AF2-multimer pipeline are likely to be suboptimal, posing significant challenges for accurately predicting interchain interactions. In contrast, our approach may be beneficial in addressing this limitation by leveraging protein-protein structural complementarity to explore potential interaction signals, thereby enhancing the prediction of interchain relationships in such biologically relevant complexes.

In **Fig. 2d**, we provide a comparative analysis of the prediction results for the eight intertwined complexes, which are characterized by their highly interwoven and tight interfaces, presenting a challenging test for accurate structure prediction. The figure highlights the average values of multimer TM-score, monomer TM-score, and interface DockQ for both DeepSCFold and NBIS-AF2-multimer. Our method demonstrates superior performance across all metrics: the multimer TM-score improves from 0.68 (NBIS-AF2-multimer) to 0.82 (DeepSCFold), the monomer TM-score increases from 0.67 to 0.77, and the interface DockQ rises from 0.45 to 0.60. To further elucidate these findings, **Fig. 2g** showcases a representative example from T1161, an intertwined complex comprising 96 residues. In this case, the structure model predicted by NBIS-AF2-multimer achieves a monomer TM-score of 0.44, which significantly compromises the overall complex modeling accuracy, yielding a complex TM-score of 0.45. In contrast, DeepSCFold achieves a monomer TM-score of 0.92, resulting in a substantially higher complex TM-score of 0.96. These results underscore the importance of structure-level information inferred from sequences, which not only enhances the determination of inter-chain distances and orientations but also complements the co-evolutionary signals derived from monomeric MSAs constructed through sequence similarity. This dual approach enables more accurate and reliable predictions of complex structures, particularly in challenging cases such as intertwined complexes.

### Structural modeling for antibody-antigen complexes with DeepSCFold

Given the inherently weak or absent co-evolutionary signals between antibody and antigen sequences, we conduct a focused investigation into the performance of antibody-antigen complex modeling, with particular emphasis on the impact of complex paired multiple sequence alignments (MSAs). We evaluated DeepSCFold on a curated set of antibody-antigen targets extracted from SAbDab, spanning entries from January 25, 2024, to June 1, 2024. This temporal separation ensures no overlap with the training data of the deep learning model or the protein database utilized for structure modeling, thereby maintaining the integrity of the evaluation. For a fair and objective comparison, both DeepSCFold and AlphaFold-Multimer were provided with identical template and monomeric MSAs. Notably, the complex paired MSA employed by DeepSCFold was constructed by deriving alignments for each subunit from their respective monomeric MSAs. Additionally, we included a comparison with the latest AlphaFold3^26^, with results obtained directly from its online server (https://alphafoldserver.com/) to ensure up-to-date benchmarking.

**Fig. 3a** illustrates the complex TM-scores and interface DockQ values of the predicted antibody-antigen complex models, benchmarked against their corresponding experimental structures in the PDB database. The interface DockQ specifically assesses the interaction between the antibody and antigen chains, excluding the interface between the antibody’s heavy and light chains. Detailed evaluation results are provided in **Supplementary Table 2**. On average, DeepSCFold achieves a complex TM-score of 0.73 and an interface DockQ of 0.39, outperforming both AlphaFold-Multimer (TM-score = 0.67, DockQ = 0.25) and AlphaFold3 (TM-score = 0.72, DockQ = 0.37). **Fig. 3b** provides a comparative analysis of prediction accuracy between DeepSCFold and AlphaFold-Multimer/AlphaFold3 for each target. DeepSCFold achieves higher TM-scores than AlphaFold3 in 59.6% of the test cases and higher DockQ scores in 60.7% of the cases. Additionally, the prediction success rate for the antibody-antigen binding interface (DockQ > 0.23) is improved by 12.4% with DeepSCFold compared to AlphaFold3. These results underscore the complementary nature of the two methods, suggesting that DeepSCFold and AlphaFold3 can synergistically enhance overall prediction performance. Compared to AlphaFold-Multimer, DeepSCFold demonstrates superior performance in 77.5% of test cases based on TM-score and 92.1% based on DockQ, highlighting its ability to effectively augment the performance of its structural prediction component, AlphaFold-Multimer, through the generation of paired MSAs. Notably, while AlphaFold-Multimer produced models with DockQ < 0.23 for 56 targets, DeepSCFold successfully built antibody-antigen complex models with DockQ > 0.23 for 39.3% of these challenging cases. This improvement suggests that the complex paired MSAs constructed based on structural complementarity provide valuable inter-chain interaction signals, thereby enhancing the accuracy of complex modeling.

**Fig. 3.**
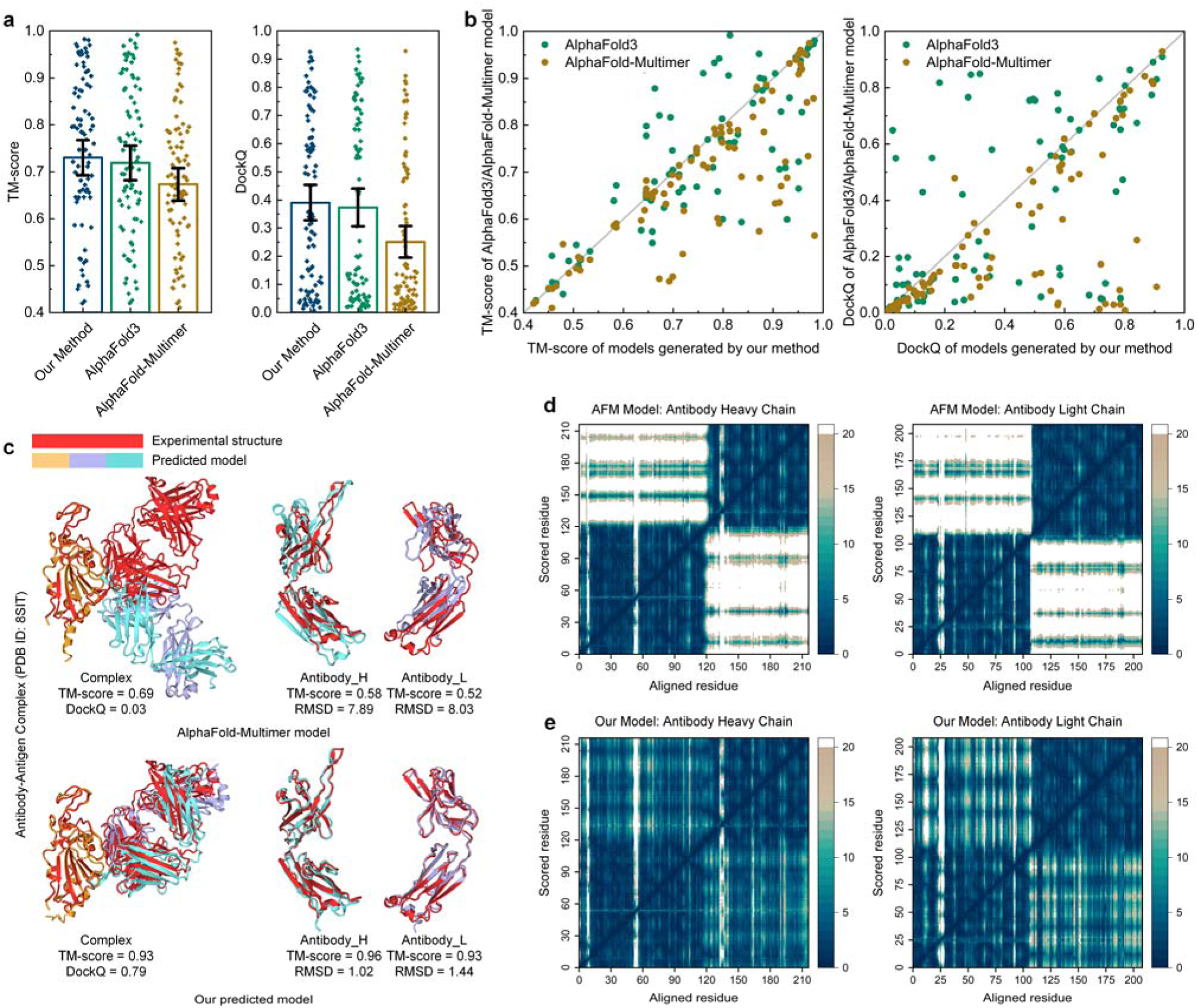
Results of antibody-antigen complex structure prediction. **a.** Average TM-scores of complex structures and DockQ scores of antibody-antigen interface for the predicted models by DeepSCFold, AlphaFold-Multimer and AlphaFold3. Error bars are 95% confidence intervals. **b.** Head-to-head comparison of the TM-scores and DockQ scores on each test case between DeepSCFold, AlphaFold-Multimer and AlphaFold3. **c.** Example of an antibody-antigen complex (PDB ID: 8SIT) predicted by DeepSCFold and AlphaFold-Multimer, where experimental structures are colored in red and monomers of the predicted models are colored differently. **d-e.** Predicted aligned error (PAE) plots for the antibody heavy and light chains modeled by AlphaFold-Multimer (**d**) and DeepSCFold (**e**).

A notable example is the complex structure of the SARS-CoV-2 spike receptor-binding domain with the broadly neutralizing antibody CC84.24 Fab (PDB ID: 8SIT). AlphaFold-Multimer’s prediction for this protein complex exhibits significant errors in inter-chain orientation, achieving an overall TM-score of 0.63 and an interface DockQ of 0.03 (**Fig. 3c**). Furthermore, AlphaFold-Multimer generated low-quality monomer models for the antibody, with the heavy chain (TM-score = 0.58, RMSD = 7.89 Å) and the light chain (TM-score = 0.52, RMSD = 8.03 Å) both showing substantial deviations from the experimental structure. We further assess the inter-domain accuracy of single-chain structure model by the predicted aligned error (PAE) plot, which captures the model’s global inter-domain structural errors^57^. The PAE plots for the antibody heavy chain and light chain structures predicted by AlphaFold-Multimer are shown in **Fig. 3d**. These plots reveal two distinct low-error squares corresponding to the two domains of each single-chain structure. While the PAE values within individual domains are low, with each domain achieving a TM-score above 0.9, the inter-domain orientations exhibit high PAE values, indicating that AlphaFold-Multimer fails to accurately predict the relative domain orientations in this case. In contrast, DeepSCFold not only accurately models each domain but also captures the correct inter-domain and inter-chain orientations, as evidenced by the low PAE values across the entire region for both the heavy and light chains (**Fig. 3e**). These results highlight that, for flexible proteins such as antibodies, while monomer MSAs can effectively infer domain-level topology, accurate structure modeling often requires complex-wide MSAs to capture the interaction signals necessary for resolving their orientations. This emphasizes the importance of incorporating structural complementarity and inter-chain interaction signals in predicting biologically relevant conformations of antibody-antigen complexes.

### The performance of sequence-based structural similarity prediction

One core component of DeepSCFold is the prediction of protein-protein structural similarity directly from sequence data. The co-evolutionary signals derived from homologous protein searches are essential for protein structure prediction, as they shed light on residue-residue interactions and evolutionary constraints that shape protein folding and complex formation. Due to the low cost and large scale of sequence data, sequence-based search methods are generally more convenient and faster than structure-based methods, which is especially evident in scenarios involving a large number of new sequences^41^, such as metagenomic sequences^53^ and antibody variant sequences^58^. Nevertheless, in cases of highly distant evolutionary relationships, sequences may have diverged so extensively that identifying their relatedness becomes challenging^59^. This challenge highlights the need for developing novel computational approaches that can directly infer protein-protein structural similarity from sequence information, thereby complementing existing structure prediction methods.

To enhance detection performance while ensuring the versatility and efficiency of sequence search, we developed a novel deep learning model that directly predicts protein structural similarity from sequences, serving as the first core component of DeepSCFold (see details in Methods). We hypothesize that structurally similar protein sequences, despite exhibiting substantial sequence divergence, can effectively address the limitations of conventional sequence alignment methods by providing crucial structural topology information, thereby establishing a robust foundation for constructing high-quality multiple sequence alignments (MSAs) of monomeric proteins, which is essential for achieving high-accuracy structural modeling. This conceptual framework holds particular significance for the alignment analysis of distantly related homologous proteins, offering new insights into the evolutionary relationships among protein families with low sequence similarity. Our predicted protein-protein structural similarity (pSS-score) is represented by the template modeling score (TM-score), which is a widely used metric for assessing the global similarity between two protein structures. For a comprehensive and objective analysis, we established a benchmark dataset comprising CASP15 target proteins, implementing a temporal cutoff to guarantee complete independence from the training data, and compared the performance of our method with the state-of-the-art method PLMsearch^41^ using Pearson^60^, Spearman^61^, and ROC (AUC)^62^ statistical metrics. As shown in **Fig. 4a**, the correlation between the pSS-score and the true TM-score (calculated by the US-align^63^ tool) for protein pairs in the dataset was measured using the Pearson and Spearman metrics. Our method achieved a Pearson correlation of 0.83 compared to 0.79 for PLMsearch, indicating a 5.06% improvement. Notably, the Spearman correlation, which evaluates the rank-order relationship, was 0.69 for our method and 16.95% higher than the 0.59 achieved by PLMsearch. This significant improvement in Spearman correlation demonstrates that our model more effectively preserves the relative ordering of structural similarities, which is crucial for applications involving the ranking of protein similarity. Regarding the receiver operating characteristic (ROC) curve analysis, which measures the model’s discriminative capability between positive and negative samples, our method demonstrated superior performance with an area under the curve (AUC) of 0.87, representing a 12.99% improvement over PLMsearch’s AUC of 0.77. For this evaluation, positive cases were rigorously defined as protein pairs exhibiting true TM-scores within the highest quartile of the dataset distribution. This further demonstrates the improved classification ability of our method in identifying high-confidence protein pairs, demonstrating robust performance in structural similarity prediction. The details of these results are provided in **Supplementary Table 3**.

**Fig. 4.**
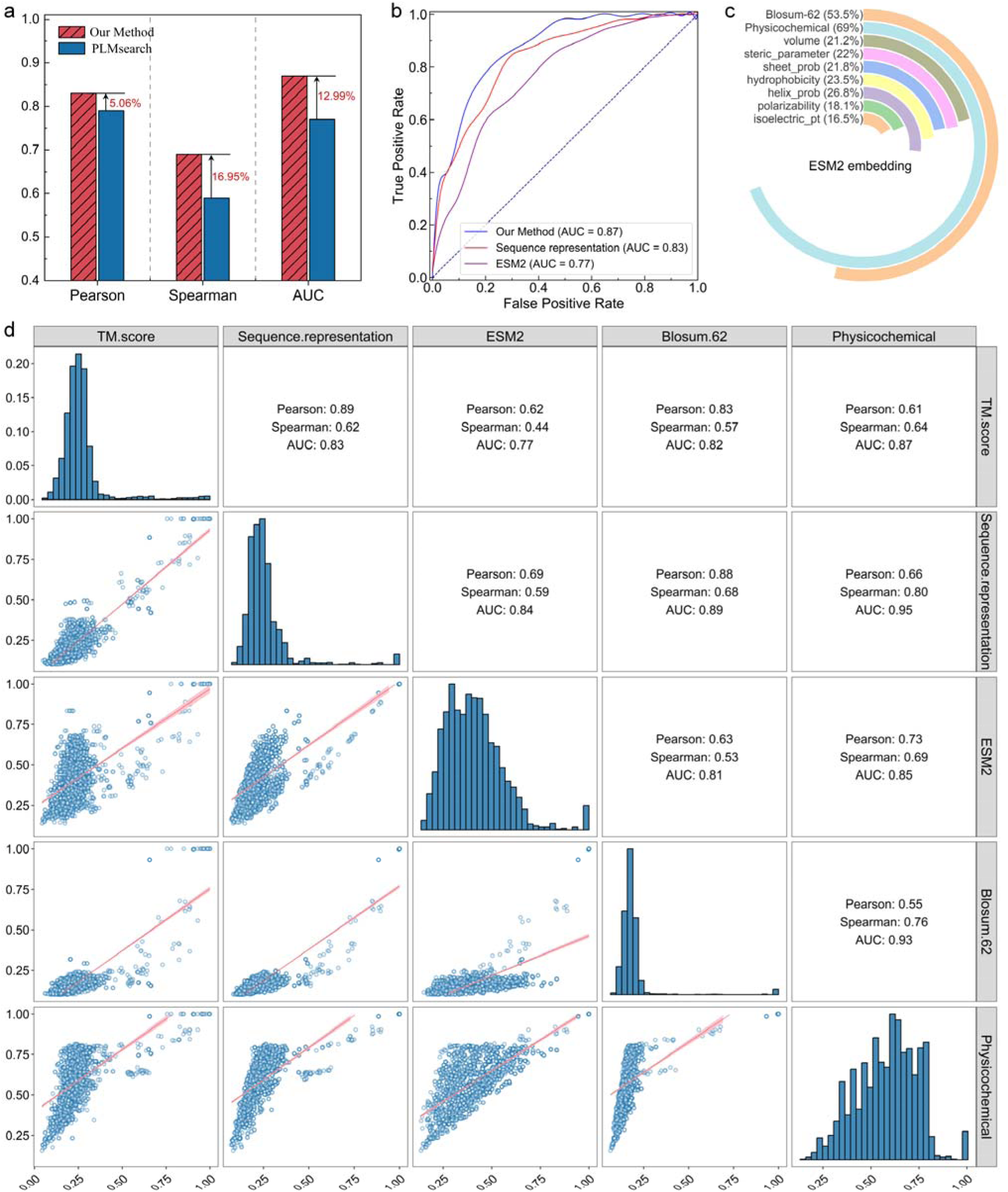
Comparative performance and feature analysis for protein-protein structural similarity prediction. **a.** Performance comparison between our method and PLMsearch based on Pearson correlation, Spearman correlation, and ROC AUC metrics. **b.** ROC curves showing the classification performance of our method, ESM2, and Sequence representation. **c.** Correlation analysis between ESM2 embeddings and additional sequence-derived features. **d.** Feature distributions and pairwise correlations. Histograms represent the distribution of each feature, scatter plots show pairwise relationships, and upper triangle statistics include Pearson, Spearman correlations, and AUC values.

The comparative analysis presented in **Fig. 4b** systematically investigates the potential factors contributing to the enhanced performance of our method through a comprehensive evaluation against ESM2 embedding and the sequence representation generated from our method. In the ESM2 method, we used the sequence embeddings generated by the ESM2 model and calculated similarity scores between these embeddings, which resulted in an AUC of 0.77. This result shows that the protein language model effectively captures the biological information encoded in the sequence, and both our method and PLMsearch use ESM-based embeddings as a foundation. Importantly, our method further combines additional sequence-derived features to generate the sequence representation, such as physicochemical properties and amino acid substitution probabilities^64^. To evaluate their impact, we extract sequence representations through the multi-scale retention module with the concatenated features, and then calculated the similarity scores using the same procedure as the ESM2 baseline. The results show that the AUC of the sequence representation reaches 0.83, indicating that the additional features, which contain biological constraints, positively contribute to the performance compared to the ESM2 embedding alone. In addition, the performance gap between sequence representation (AUC = 0.83) and our full method (AUC = 0.87) demonstrates the importance of our classification module, which effectively utilizes the combined features to achieve more reliable classification and ranking performance.

In **Fig. 4c**, we conducted a comprehensive analysis of the correlation between ESM2 embeddings and various sequence-derived features to evaluate their potential complementarity. The circular diagram illustrates these relationships, where each arc corresponds to a distinct feature, with the accompanying percentage quantitatively representing its correlation with the ESM embedding. The overall physicochemical feature set exhibited the highest correlation with ESM2 embeddings (69.0%), indicating some overlap in the biological information captured by both. However, individual properties such as hydrophobicity (23.5%), helix probability (26.8%), and polarizability (18.1%) showed relatively low correlations, suggesting that each property contributes unique information not fully captured by the ESM2 model. Notably, the amino acid substitution matrix (Blosum-62) shows only a 53.5% correlation with ESM2 embeddings, demonstrating that evolutionary information encoded by substitution probabilities provides complementary insights.

Specifically, we present a comprehensive overview of the distribution and correlation among various features and their relationship with true TM-score (**Fig. 4d**). This experimental design enables the identification of key components driving the performance improvement while providing insights into the relative contributions of different sequence features. The figure consists of histograms along the diagonal, illustrating the distribution of values for each individual item: TM-score, sequence representation, ESM2 embeddings, Blosum-62, and physicochemical properties. The sequence representation exhibited the highest correlation with TM-score (Pearson’s r = 0.89, Spearman’s ρ = 0.62), demonstrating the effectiveness of integrating diverse biological information. In contrast, ESM2 embeddings alone showed lower correlations (Pearson’s r = 0.62, Spearman’s ρ = 0.44), suggesting that ESM2 captures essential biological information but benefits from further enhancement through additional features. The Blosum-62 matrix demonstrated a Pearson correlation of 0.83 and a Spearman correlation of 0.57 with TM-score, indicating the contribution of evolutionary information to structural similarity. Physicochemical features also showed a strong relationship with structural similarity, with a Spearman correlation of 0.64 and an AUC of 0.87. Furthermore, the correlations between different features varied, indicating that while there was some overlap, each feature set provided unique information. These results demonstrate the necessity of integrating diverse features, as this combination allows the model to capture a more comprehensive representation of biological characteristics, thereby improving its ability to predict sequence-to-structure relationships.

### The performance of protein-protein interaction probability from sequences

DeepSCFold directly predicts protein-protein interaction (PPI) probabilities through sequence data analysis as another key capability. Many existing methods for constructing protein paired MSAs rely on experimental data and manually annotated information, such as species-specific annotations. This reliance may limit their applicability, particularly when dealing with less-studied or unknown species. If the interaction probability between two given sequences could be directly predicted, it would greatly expand the data sources available for MSA construction. This approach would allow the use of large-scale, unannotated protein sequence databases, such as the BFD^51,52^ and MGnify^53–55^ metagenomic databases, which have been shown to contribute significantly to protein structure prediction.

To evaluate the performance of our method on interaction prediction, we trained our model on a dataset consisting of human protein pairs, and compared the results with those of state-of-the-art PPI prediction methods, Topsy-Turvy^65^ and RAPPID^66^, across four distinct species: *Saccharomyces cerevisiae* (Yeast), Escherichia coli (E. coli), Drosophila melanogaster (Fly), and Caenorhabditis elegans (Worm). **Fig. 5a** illustrates the F1-scores of our method compared to Topsy-Turvy and RAPPPID across all test species, including an aggregated result (“ALL” in **Fig. 5a**). Our method consistently achieved higher F1-scores, indicating its strong ability to balance precision and recall, as well as its robustness in predicting interactions within diverse biological systems. Notably, the model performed better in Drosophila and C. elegans, two organisms known for their intricate PPI networks due to complex developmental processes and signaling pathways. These results indicate that our method has more robust cross-species generalization performance and effectively captures potential co-evolutionary signals. The details of results are provided in **Supplementary Table 4**.

**Fig. 5.**
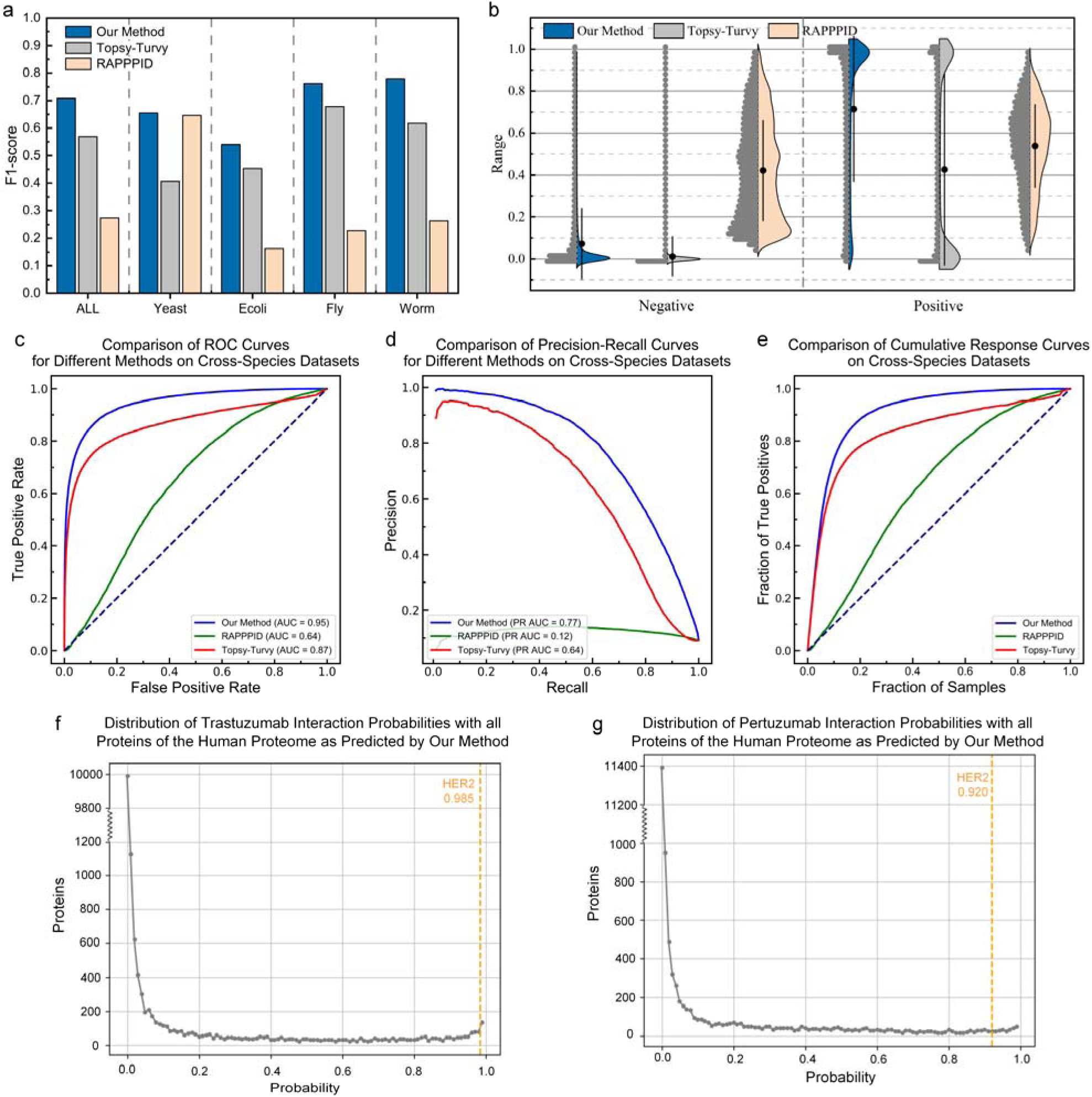
Comparative analysis and practical applications of protein-protein interaction (PPI) prediction. **a.** F1-scores of our method compared to SOTA approach Topsy-Turvy and RAPPPID across different species (Yeast, E. coli, Fly, Worm) and the aggregated dataset (ALL). **b.** Violin plots showing the distribution of interaction probability scores for positive and negative protein pairs across methods. **c-e.** Receiver operating characteristic (ROC) curve, Precision-recall (PR) curve, and cumulative response curve (CRC) analysis of our method compared to Topsy-Turvy and RAPPPID. **f-g.** Distribution of interaction probabilities of Trastuzumab (**f**) and Pertuzumab (**g**) with the human proteome, highlighting high interaction scores with HER2 (Trastuzumab: 0.985, Pertuzumab: 0.920).

The violin plots in **Fig. 5b** display the distributions of interaction probability scores for both negative and positive protein pairs across all test datasets. For each method, the plot indicates the spread, median, and overall distribution of predicted scores. For negative samples, our method shows a narrow distribution close to zero, highlighting its ability to accurately classify non-interacting pairs with a low false negative rate. However, the most significant observation is seen with positive samples. Moreover, our method exhibits a concentrated distribution of probabilities above 0.8 for positive samples, indicating high certainty in correctly identifying the interaction pairs. In contrast, the distribution of Topsy-Turvy is divided into two regions, one with high confidence and the other with low confidence, revealing the limitations of its predictions. The results of RAPPPID showed that the distributions were not well-separated, leading to low overall discrimination performance. In summary, our method shows better performance in distinguishing interaction pairs, with potential to identify structural complementarity between proteins.

To further assess the robustness of DeepSCFold in predicting protein-protein interactions (PPIs), we employed three standard evaluation metrics: the receiver operating characteristic (ROC) curve, the precision-recall (PR) curve, and the cumulative response curve (CRC). The ROC curve analysis compares the true positive rate (sensitivity) to the false positive rate across different decision thresholds (**Fig. 5c**). DeepSCFold achieved an area under the curve (AUC) of 0.95, outperforming RAPPPID (AUC = 0.64) and Topsy-Turvy (AUC = 0.87). The higher AUC indicates that DeepSCFold consistently performs well in distinguishing true interactions from non-interactions across a wide range of confidence thresholds. The PR curve analysis (**Fig. 5d**) provides insight into model performance under imbalanced conditions, where the positive class (interacting protein pairs) is less frequent, as is typical in real-world PPI datasets. The PR AUC for DeepSCFold was 0.77, outperforming RAPPPID (0.12) and Topsy-Turvy (0.64). This higher PR AUC reflects DeepSCFold’s ability to maintain high precision without sacrificing recall, which is essential for minimizing false positives while retaining a large proportion of true interactions. The cumulative response curve in **Fig. 5e** illustrates the fraction of true positive PPIs identified as a function of the sample size evaluated. The curve of DeepSCFold rises more steeply than those of Topsy-Turvy and RAPPPID, indicating that it can capture a considerable proportion of true interactions with fewer samples. This rapid enrichment of true positives makes DeepSCFold effective for identifying high-confidence PPIs early in exploratory studies, thereby accelerating the discovery of potential interactions.

To demonstrate the practical utility of DeepSCFold, we performed an in-depth analysis of the interaction probabilities of two well-studied antibodies, Trastuzumab and Pertuzumab^67^, with the entire human proteome (17,286 proteins from the STRING^68^ database). The distribution of predicted interaction scores (**Fig. 5f** for Trastuzumab and **Fig. 5g** for Pertuzumab) confirms that the model assigns extremely high interaction probabilities to known interactions with HER2, a critical receptor targeted by both antibodies. The results for Trastuzumab showed an interaction score of 0.985 with HER2, while Pertuzumab’s score was 0.920, aligning with biological evidence of their binding specificity. The sharp distribution of interaction probabilities in these analyses highlights DeepSCFold’s potential in large-scale interaction screening, paving the way for more efficient drug discovery and functional annotation of proteins.

## Discussion

Accurately capturing inter-chain orientations and determining the structure of protein complexes remains a significant challenge, as complex structure prediction is less accurate than single-chain modeling. In this work, we have proposed DeepSCFold, a protocol for automated protein complex prediction that employs deep learning networks to capture potential structural complementarity and construct complex MSAs. The results demonstrate that DeepSCFold could provide interaction signals that effectively improve the structural prediction accuracy of AlphaFold-Multimer.

It has been reported that the ways of protein-protein interactions in nature may be limited, with similar binding modes can be identified for almost all known protein-protein interactions. To capture interaction signals indicative of protein-protein binding modes, DeepSCFold employs deep learning models to predict protein structural similarity and interaction probabilities, which are subsequently used to construct complex MSAs. Notably, current sequence concatenation mechanisms based on species annotations are limited to genomic sequences and thus cannot fully utilize the highly informative homologous sequences available in metagenomic databases to guide multi-chain structure assembly. DeepSCFold avoids this limitation and provides a new possibly way for the monomeric MSAs concatenation. By integrating complex MSAs with the state-of-the-art AlphaFold-Multimer modeling method, DeepSCFold enhances the accuracy of complex structure prediction, as borne out by experimental results on the CASP15 protein complex dataset.

Antibody-antigen complexes are widely recognized as lacking inter-chain co-evolution, as it is not possible to find orthologs between host and pathogenic proteins as there is no species overlap (e.g. humans and a virus, originating from different kingdoms). However, when applied to antibody-antigen complex modeling, DeepSCFold achieved a 24.7% higher success rate in predicting antibody-antigen interfaces (DockQ > 0.23) compared to AlphaFold-Multimer, using the same monomeric MSAs and templates. This improvement provides further evidence supporting our hypothesis that the enhanced performance stems from DeepSCFold’s ability to capture shape and physicochemical complementarity through its structure-aware complex MSA construction, overcoming the inherent limitations of inter-chain co-evolution in antibody-antigen systems.

Despite the improved performance in complex modeling, there are still some challenges for DeepSCFold. Users are required to provide stoichiometric information for the complex (e.g., the copy number of each component chain) prior to executing DeepSCFold, which may limit its practical applicability. Accurate model quality assessment (MQA) is crucial for reliably distinguishing high-accuracy predictions from incorrect ones, especially in the absence of experimental structures for validation. Moreover, MQA provides users with a self-evaluation confidence score for the model, offering valuable insights into the reliability of the predicted models. Extending DeepSCFold beyond protein-protein complexes to simulate structures formed by interactions between DNA, RNA, and other ligands and proteins is also a topic to explore in our ongoing research.

## Methods

### Training set

For training a sequence-based structural similarity prediction model, we collected protein complex structures from the Protein Data Bank (PDB, https://www.rcsb.org/) up until June 2022 and applied the following filtering criteria: (1) resolution better than 3.5 Å; (2) with the length of each chain greater than 50 residues; (3) the number of chains in the complex does not exceed three; and (4) the complex structure does not contain DNA or RNA. After filtering, a total of 36,955 protein complexes were retained. Next, we employed DomainParser^69^ to split the protein chains in these complexes into domains, and then clustered the resulting candidate domains using MMseqs2^36^ with a 30% sequence identity threshold. After this step, we obtained 15,636 non-redundant domains. To assess the structural similarity between these domains, we performed pairwise comparisons of the non-redundant domains using USalign^63^ and calculated the TM-score, which ranges from 0 to 1, with higher scores indicating greater structural similarity. To address the potential bias in the data distribution, we divided the TM-scores into 10 bins, each spanning a 0.1 interval, ensuring that each bin contained an equal number of domain pairs. This process resulted in approximately 100,000 domain pairs for subsequent training.

For training a sequence-based interaction probability prediction model, we extracted positive examples from the STRING^68,70^ database (v11.5) based on physical binding interactions associated with high experimental evidence scores. From this set, we removed protein pairs involving very short proteins (shorter than 50 amino acids) and, due to GPU memory constraints, also excluded proteins longer than 1000 amino acids. Next, we removed protein-protein interactions with high sequence redundancy from the dataset to prevent the model from memorizing interactions based on sequence similarity alone. Specifically, we clustered the proteins at a 40% sequence identity threshold using MMseqs2^36^. A PPI (A-B) was considered redundant and excluded if another PPI (C-D) had already been selected, where the protein pairs (A, C) and (B, D) were both found within the same clusters. After this step, we obtained 39,183 positive samples. Non-interactions (‘negatives’) were sampled randomly from the respective proteins in the positive set, with a positive-to-negative sample ratio of 1:10.

### Test set

To assess the performance of protein complex prediction, a total of 25 multimer targets were selected from the CASP15 dataset, as listed under the MULTIMERS category on the official CASP website (https://predictioncenter.org/casp15/). These targets were chosen based on their suitability for structural modeling using AlphaFold-Multimer on a single NVIDIA A100 GPU, which is widely used for large-scale protein structure predictions. All selected targets have experimentally determined structures available in the Protein Data Bank (PDB), providing a reliable benchmark for comparison. To evaluate the performance of DeepSCFold on protein complexes lacking inter-chain co-evolution information, we further constructed an independent test set of antibody-antigen protein complexes. We collected antibody-antigen complexes with antigen types classified as protein from the SAbDab^48^ database, spanning the period from January 25, 2024, to June 1, 2024. The complexes included in the dataset had a sequence length of fewer than 2000 amino acids and a resolution <5 Å. For complex entries sharing the same protein ID, clustering was performed using MMseqs2^36^ with a sequence identity threshold of 100%, leading to a final set of 89 Antibody-Antigen protein complexes.

To objectively evaluate the performance of structural similarity prediction model, we constructed a test dataset with a temporal cutoff to ensure its independence from the training data. We split the protein targets from CASP15 into individual chains and then clustered the candidate chains using MMseqs2 with a 100% sequence identity threshold, resulting in a final set of 74 unique single-chain proteins. Finally, USalign was employed to perform pairwise comparisons and calculate TM-scores, resulting in a total of 2,700 protein pairs. Furthermore, we evaluated the performance of DeepSCFold in predicting interaction probabilities using a publicly available interaction dataset from a previous study^65,68^. The dataset includes data from four species: *Drosophila melanogaster*, *Caenorhabditis elegans*, *Saccharomyces cerevisiae*, and *Escherichia coli*. For each of the four model organisms, there are 5,000 positive and 50,000 negative interactions, except for *Escherichia coli* (2,000/20,000), resulting in a 1:10 positive-to-negative ratio.

### Structural similarity and interaction probability prediction

Paired-MSA is crucial for protein multimer structure modeling. However, the construction method that only uses traditional sequence similarity combined with existing complex structure information may still face difficulties in providing high-quality paired MSAs. To efficiently construct paired MSAs, we developed a sequence-based deep learning model to capture the relationships between sequences. The process begins by extracting sequence-derived features and integrating them with embeddings obtained from a protein language model (PLM). Next, the combined features of the two query sequences are independently encoded into sequence representations through a multi-scale retention module. To capture inter-sequence relationships, these representations are then processed by a cross-attention module, which generates a sequence pair representation. Finally, this pair representation is passed through a down-sampling module to predict the structural similarity score (pSS-score) or the interaction probability score (pIA-score).

### Model architecture and input features

We develop a deep learning model to predict the inter-chain interaction and the structure similarity from the input sequences. The input features for the network encompass a diverse set of information, including residue type, positional information, evolutionary relationships, physicochemical properties, and contextualized embeddings. Specifically, these features include one-hot encoding, normalized distance to sequence end, Blosum-62 substitution matrix, physicochemical features^64^, and embeddings from the pre-trained ESM2^40^ language model. Together, these features provide a comprehensive representation of each residue, capturing sequence, structural, and functional properties. Detailed descriptions of all input features are provided in **Supplementary Table 5**.

The network architecture consists of a multi-scale retention module, a criss-cross attention module, and a down-sample module. Initially, the input query protein sequence features are processed and fused to generate task-specific sequence representations. The ESM2 embeddings are first passed through a linear layer for dimensionality reduction, followed by ReLU activation and LayerNorm to introduce non-linearity and normalize the feature distributions. This ensures that the embedding dimensions are appropriately aligned with the 1D features, preventing a large dimensional gap that could otherwise obscure the contribution of the 1D features. The reduced ESM2 embeddings are then concatenated with the 1D input features to form a combined feature embedding. Subsequently, the output is fed into a multi-scale retention module to generate a sequence representation by capturing long-range dependencies. This module consists of 8 layers, with each layer combining a multi-scale retention mechanism and a feed-forward network to refine the information. Next, the relationships between the proteins are considered. The sequence representations of the two queries are combined through matrix multiplication, followed by Conv2D and ELU activation to generate a combined sequence representation. This representation is then processed through the criss-cross attention module that captures interactions along both the horizontal and vertical axes to generate a sequence pair representation. The module computes separate attention maps for each direction using the query, key, and value projections, followed by softmax normalization to generate attention weights. These attention weights are applied to the value tensors, producing an updated sequence pair representation that captures the relationships between the query proteins. Finally, the sequence pair representation is fed into the down-sample module, which consists of four down-sample blocks and four residual blocks. Each down-sample block is composed of a LayerNorm followed by a convolutional layer, with the first block using a 4×4 convolution and the other three blocks using 2×2 convolutions. The residual blocks are structured with a series of operations: a 7×7 convolution, LayerNorm, a 1×1 convolution, GELU activation, and a second 1×1 convolution. The output of the down-sample module is passed through a fully connected layer, followed by a sigmoid function to generate the final prediction scores.

### Training

During the training, 90% of the protein pairs are randomly selected from the development set as the training set, and the remaining 10% as the validation set. The network is implemented in Python using PyTorch and trained for up to 30 epochs. Due to GPU memory constraints, the training batch size starts at 4 and is progressively reduced as training progresses: it is set to 4 for the first 10 epochs, reduced to 2 for the next 10 epochs, and finally lowered to 1 for the last 10 epochs. The Adam optimizer is employed to minimize the loss, with the total loss calculated using the Log-Cosh Loss function for the structural similarity prediction model and the Focal Loss function for the protein-protein interaction prediction model. The learning rate is initially set to 1e-5 and decreases progressively during training, following an exponential decay strategy. The network model with the least validation loss is selected for downstream tasks. It takes about 14 days to train the network using one NVIDIA Tesla V100 GPU and 80 CPU cores.

### MSA sampling and paired-MSA construction

In order to obtain more diverse MSAs, we first search multiple sequence databases (UniRef30^49^, UniRef90^50^, UniProt^50^, BFD^51,52^, MGnify^53–55^, and the ColabFold DB^56^) to obtain the MSAs of each monomer. The detailed descriptions of above databases are given in **Supplementary Note 2**. Then, we use the predicted pSS-score as a supplement to the traditional sequence similarity in the ranking and selection of the searched MSAs. For multimers, we use the developed deep learning model to predict the pIA-scores for each pair of sequence alignments from different subunit MSAs. The subunit MSAs are then concatenated based on these interaction probabilities to construct paired MSAs. Additionally, we use information from multiple sources, such as species annotations, UniProt accession number, and protein complexes from the Protein Data Bank (PDB), to further construct a series of paired MSAs. The paired MSAs constructed above are used in the structure predictor to enhance diversity and prevent falling into local optimum.

### Quality assessment of built models

In this study, we employed an in-house developed single model quality assessment method based on multi-scale protein representations using hierarchical networks. To represent the protein, physical and geometric properties of residues are extracted at the 1D scale, interaction relationships at the 2D scale, and structural topology at the 3D scale. These multi-scale representations are input into a multi-layer network module, which includes graph attention, transformer, and convolutional networks, to predict model quality. Building on this scoring model, we further developed a multi-model evaluation method based on structural consensus. First, the method selects the top N predicted models as the high-quality model pool using the single model quality assessment and model self-assessment scores. Next, all predicted models are aligned with the high-quality model pool to calculate their respective quality scores, and the top 5 models are selected for final evaluation.

### Evaluation metrics

We use two types of metrics to evaluate the quality of the protein complex model built by DeepSCFold. The first type of metrics is one that measure the closeness between the built model and the experimental structure in PDB. In this respect, we adopt the TM-score^71^ between the predicted complex model and the corresponding experimental structure calculated by TM-score tool downloaded at https://zhanggroup.org/TM-score/. The second type is one that measures the interface quality of predicted protein complex model. In this regard, we report DockQ^72^ score calculated by public tool (https://github.com/bjornwallner/DockQ/). Besides the above metrics, we also reported the Pearson, Spearman and ROC-AUC/PR-AUC to evaluate the performance of protein-protein structural similarity and interaction probability prediction models. The detailed descriptions of above metrics are given in **Supplementary Note 1**.

### Statistics and reproducibility

All data were carefully collected and analyzed using standard statistical methods. Comprehensive information on the statistical analyses used is included in various places, including the figures, figure legends and results.

### Reporting summary

Further information on research design is available in the Nature Portfolio Reporting Summary linked to this article.

## Data availability

The authors declare that the data supporting the results and conclusions of this study are available within the paper and its Supplementary Information. The CASP15 data including the experimental structures are available at: https://predictioncenter.org/casp15/index.cgi. The antibody-antigen data from SAbDab database is available at: https://opig.stats.ox.ac.uk/webapps/sabdab-sabpred/sabdab.

## Code availability

The online server of DeepSCFold is made freely available at http://zhanglab-bioinf.com/DeepSCFold/.

## Acknowledgements

We thank members of the Guijun Zhang lab for discussion and feedback. Computational resources were provided by the College of Information Engineering at Zhejiang University of Technology. This work was supported by the National Key R&D Program of China [2022ZD0115103], the National Nature Science Foundation of China (62173304, 62203389), the “ Pioneer ” and “ Leading Goose ” R&D Program of Zhejiang(2025C01190), the Zhejiang Province High-level Talent Special Support Program (2023R5248), and Fundamental Research Funds for the Provincial Universities of Zhejiang (RF-C2024006).

## Author contributions

G.Z. conceived the project. J.Z. helped supervise the research. G.Z., M.H., and Y.X. designed the experiment. M.H., Y.X., P.W., Z.L., and D.H. performed the experiment and collected the data. G.Z., J.Z., X.Z., M.H., and P.W. analyzed the data. M.H., Y.X. G.Z., J.Z., and X.Z. wrote the manuscript, and all authors read and approved the final manuscript.

## Additional information

**Correspondence** and requests for materials should be addressed to Guijun Zhang and Jianyang Zeng.

